# Constraining the global niche suitability of the Eusuchia clade across the Cretaceous-Paleogene boundary

**DOI:** 10.1101/2022.12.04.517697

**Authors:** Morgan Harper, Alexander Farnsworth, Paul J. Valdes, Paul J. Markwick, Maximilian T. Stockdale

## Abstract

The crocodiles and their close relatives, the alligators and gharials, have a compelling evolutionary history. They are a clade of great antiquity, with their most recent common ancestor emerging within the Mesozoic. However, unlike many groups of such a great age, the crocodilians have an extensive crown-group, with around two dozen extant examples. They have a limited ecomorphology, which has varied little since their inception, and their biogeography has been shown to interact closely with climate. The biogeography of crocodilians in deep time remains an outstanding question, which is complicated further by the limitations of the fossil record. The fossil record is fundamentally incomplete yet represents the most common method used to infer biogeography of organisms. The scarcity of fossil remains makes apparent absences difficult to confirm. Preservation bias will promote fossil occurrences in areas with a high sedimentation rate, which may not be the true ecological niche for a given taxon. This study uses species distribution models of extant crocodilians to infer the ecological niche of related taxa in the Maastrichtian and Danian. Models indicate a much wider latitudinal range than is observed among extant examples, and the invasion of new ecospace following the end-Cretaceous mass extinction. In addition, we find that while temperature is of significance to crocodilian biogeography, it is precipitation that is the most influential climatic variable.

## Introduction

Living crocodilians, comprising of crocodiles, alligators and gharials, are a very limited group both in terms of their diversity and their geographic distribution. The group contains only two dozen or so species, all of them exclusively amphibious ambush predators [1]. This lack of diversity is surprising given how ancient the crocodilians are. The common ancestor of modern crocodilians diverged from the basal Neosuchia during the early Cretaceous period [1–2]. Other clades of similar antiquity have achieved great species richness and morphological variation. For example, birds and squamates have achieved a diversity of many thousands of species since their inception in the Mesozoic. Crocodilians are also confined to a limited geographic range, restricted to within 30° of the equator [1].

The restricted species richness, ecomorphological variation and geographic range of living crocodilians contrasts with fossil examples. The crocodilians belong to a wider clade, the Eusuchia, which has a significant species richness of over 100 species [2]. Fossil eusuchians exhibit a greater ecomorphological variation than their extant counterparts, including terrestrial examples [4–5], and examples which achieved formidable sizes [6]. Fossil eusuchians also occupy a greater geographic range, with specimens occurring far outside the range of extant species; Eusuchian fossils have been recovered from locations as far north as the United Kingdom and Scandinavia [7]. The body plan of the crocodilians has remained largely unchanged since the Mesozoic [8], and therefore the niche suitability of extinct Mesozoic and Cenozoic Eusuchia [9], and the broader Crocodyliform clade [10] is widely suspected to also be dependent on environmental temperature [9].

The causes for the loss of species richness, morphological diversity and geographic range since the Cretaceous remains an outstanding question. The observed link between crocodile biogeography and climate [2] suggests that the slow decline in temperature through the Cenozoic may be a plausible culprit. Living Eusuchia cannot generate their own body heat, and therefore the impact of environmental temperature on their biogeography is to be expected. However, temperatures in the earliest stages of the Cenozoic are thought to have been similar to the Cretaceous period [11]. Analysis of ecomorphology in Eusuchians has observed a drop in diversity across the Cretaceous-Palaeogene boundary [12], seemingly prior to the Cenozoic decline in temperature. The abundance of crocodiles has been linked to distance from standing bodies of water [9], and this may have been a factor in the loss of Eusuchian diversity since the Cretaceous. Changes in the availability of standing bodies of water may have arisen due to changes in precipitation patterns or the decline in sea levels observed through the Cenozoic. The decline in eusuchian diversity and range may also be connected with the Chixulub bolide impact, widely implicated in the extinction of contemporary animal taxa such as dinosaurs [13]. It seems likely that widespread ecosystem collapse associated with mass extinction would impact the diversity and range of eusuchians. However, the species richness, morphological variation and geographic range of the Eusuchians does not appear to have recovered after the Cretaceous-Palaeogene boundary, unlike other reptile groups such as lizards [13] snakes [15].

General circulation models of palaeoclimate have diversified the portfolio of methods available to palaeontologists in recent years. Ecological niche models (ENMs), where environmental variables are used to predict biogeography, has been applied to fossil taxa in previous studies [13, 16]. These models are semi-supervised classifiers that sample maps of environmental variables using geographic coordinates of observations. Maximum Entropy classifiers have been shown to be robust predictors of ecological niches [17]. MaxEnt methodologies have also been used to construct the niche suitability of extinct biota, including Ordovician invertebrates [18], non-avian Dinosaurs [13], and Cenozoic Mammoths [19]. Preservation and observation bias makes species distribution modelling with fossils challenging, potentially skewing such models towards environmental variables that promote preservation. However, if the species being examined has a modern proxy with a comparable dependency on the abiotic environment, ENMs can be derived using occurrences of extant species, and then fitted to palaeoclimate data [16].While eusuchians certainly have significant fossil diversity, there are living examples that can be called upon to aid in the training of a species distribution model.

The analysis presented here will evaluate the role of climate in the biogeography of Eusuchians across the Cretaceous-Palaeogene boundary. Ecological niche models were used to estimate the suitable ecospace, or the ‘fundamental niche’, available to Eusuchians during the Maastrichtian and Danian. The fundamental niches of the Maastrichtian and Danian stages were compared, showing how hospitable the climate was through the Cretaceous-Palaeogene boundary, and therefore suggesting what role climate played in the decline of eusuchians. The efficacy of the ENM was tested using fossil data, demonstrating how the fundamental niche of fossil eusuchians compares with that of living examples and making inferences about their physiology and evolution.

## Materials And Methods

Modern Eusuchia species are limited to warm climates close to the equator. Analysis of the distribution of fossil occurrences has indicated that extinct members of the Eusuchia share a similar climate dependency [9]. This seems consistent with the limited ecomorphology of extinct eusuchians, which shows only limited variation compared to modern forms. This analysis used a dataset of occurrence observations of living eusuchians as a proxy for their fossil counterparts (supplementary table 1). A dataset of occurrences was download from the Global Biodiversity Information Facility (GBIF; https://www.gbif.org/) on the 20^th^ of January 2021. A comprehensive selection off search terms were used, including Eusuchia, Crocodylia, Crocodylidae, Alligatoridae and Gavialidae. GBIF is an open-source species occurrence database that has been used in previous ENM’s [20]. The GBIF occurrence data required filtering to ensure accurate spatial data was included in the model [20]. All occurrences were restricted to those of human observation. Machine observations, material samples, preserved specimens and those in zoos were excluded. All occurrence records were filtered to after 1990, to account for environmental changes and improvements in global positioning technology. A visual inspection of the occurrence data was then completed to remove any outliers, for example, those georeferenced to desert interiors, or clearly outside of viable natural locations, for example in the United Kingdom.

Occurrences with invalid or missing coordinate precision, coordinate uncertainty or occurrence date were excluded. This list is not exhaustive, and a more robust approach to evaluating GBIF spatial errors has been highlighted in Sillero and Barbosa (2020[50]). However, the data resolution criteria specified by Sillero and Barbosa is not available in the fossil data used in the later stages of this analysis. Providing all other criteria were met, entries with blank species names were retained to reflect all extant Crocodylia species belonging to the Eusuchia clade. The Crocodylidae, Alligatoridae, and Gavialidae data sets were then compiled and duplicate occurrences, identified by their occurrence number, were removed to reduce potential autocorrelation. Occurrences with the same latitude/longitude were retained providing they were distinct occurrences. Finally, the remaining occurrence data were filtered to ensure coordinate uncertainty was less than or equal to the spatial resolution of the environmental data described in section 3.3. This ensured that the recorded occurrences were directly related to the climate data in that spatial location [21]. This resulted a sample size of 14,959 Eusuchia occurrences.

ENMs require climate data as raster-formatted layers. The ENM trained in this analysis required climate data representative of the present day. This data was obtained from WorldClim (worldclim.org) v2.1 on the 5^th^ of January 2021 at a 5km spatial resolution. WorldClim bioclimatic variables represent 19 climatic variables calculated from monthly temperature and precipitation data representative of Earth’s climate between 1970 and 2000 [22]. The model was trained across an unconstrained, global extent, as recommended by Barve et al [23]. This global extent is used for throughout, removing the need to extrapolate the model to a different geospatial extent, reducing unnecessary uncertainties [24].

Maximum entropy classifiers assume mutually independent climate variables. Not all the climate variables acquired from WorldClim satisfy this criteria; for example, total annual precipitation is not independent of precipitation in the wettest quarter. Therefore it was necessary to identify a suitable selection of climate variables. The 19 variables were each compared in a pairwise manner using a pearson correlation coefficient. These correlation coefficients were expressed as Euclidean distances, and plotted on a dendrogram (Fig 1). This dendrogram was used to select the most mutually independent variables available. The variables selected were bioclimatic variables 4, 9, 10, 16, 17 and 19. These correspond to the standard deviation of temperature, the mean temperature of the driest quarter, the mean temperature of the warmest quarter, the total precipitation of the wettest quarter, the total precipitation of the driest quarter and the total precipitation of the coldest quarter.

**Fig 1.**
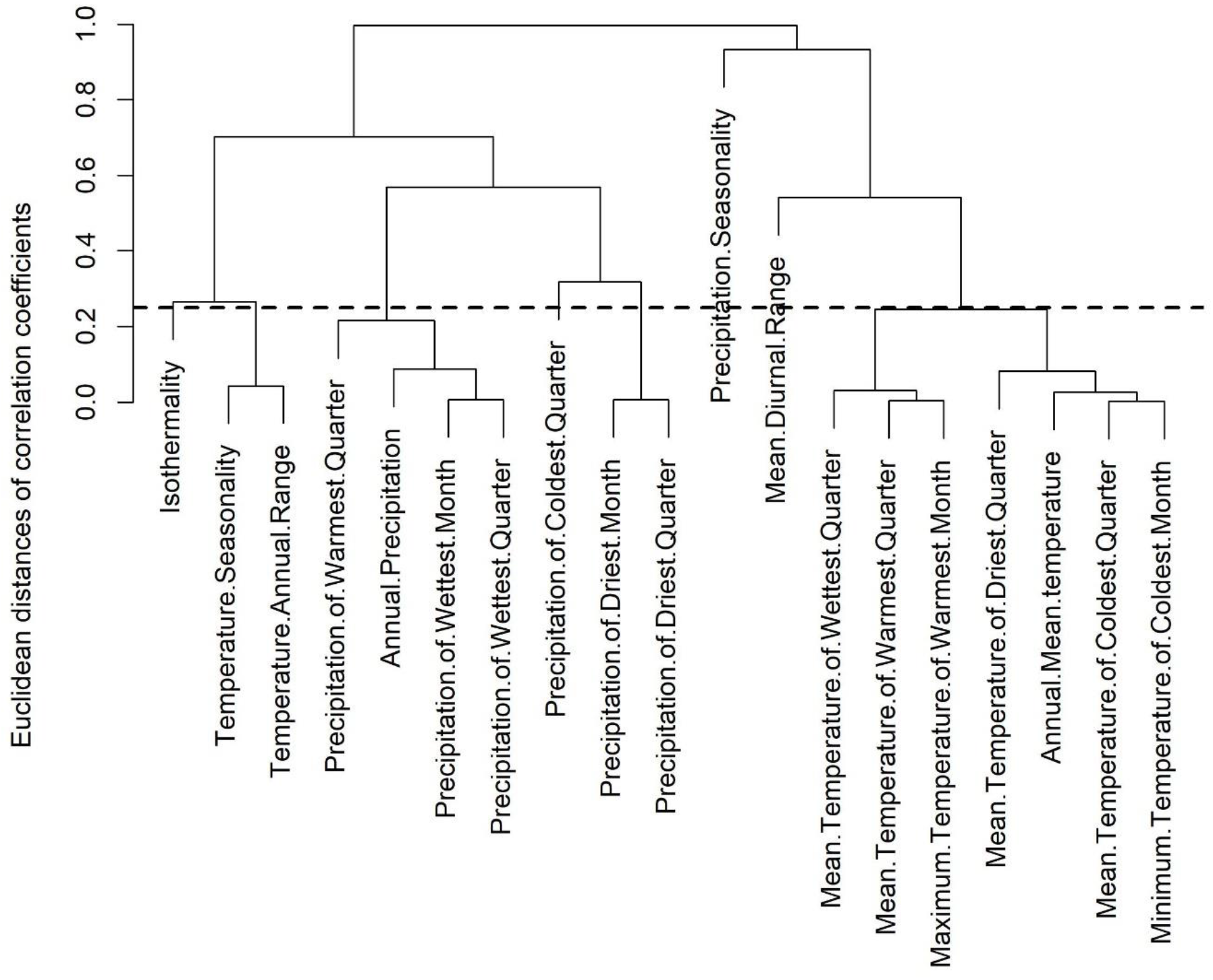
Cluster dendrogram showing the relative independence of climate variables in this analysis. Variables sharing nodes below the threshold of 0.25 were deemed mutually co-dependent, so only one variable per such node was used.

All ENMs were produced in R version (v) 3.6.0 using MaxEnt v3.4.4 and the Dismo package v1.1-4 [25]. The use of MaxEnt is considered robust due to its ability to use either absence or pseudo-absence occurrence data [13, 16]. This study used a pseudo-absence approach due to a lack of robust Eusuchia absence data, following the methodology of Waterson et al [16]. The results from MaxEnt projections using pseudo-absence data have been found to be as robust as those from true absence models [26].

Pseudo-absence MaxEnt approaches requires synthetic background noise data in place of real absence data [27]. The optimal number of pseudo-absence occurrences varies between 1000 [28] and over 10,000 [25], or can be represented as a presence: pseudo-absence ratio [29]. This analysis used an intermediate value of 5000 pseudo absence points distributed worldwide (supplementary table 2). The presence and pseudo-absence data was partitioned into test and training datasets at random, with a size ratio of 3 test points for every 7 training points. A bootstrapping procedure was employed to mitigate the impact of outlying taxa. The test and training datasets were resampled 50 times using a random 50% of rows. The ENM was implemented multiple times, once for each of the 50 resampled datasets. The outputs of each replicate were averaged to give an overall estimated ENM. This methodology yielded a worldwide map of relative niche suitability (Fig 2). The standard deviation of the data replicates yielded an additional map of model uncertainty (Fig 3). The map of niche suitability was thresholded to estimate areas where extant eusuchians occur. The chosen threshold was a relative niche probability of 0.56; this value marks the niche probability where the false negative rate and false positive rate of the maximum entropy classifier are in balance.

**Fig 2.**
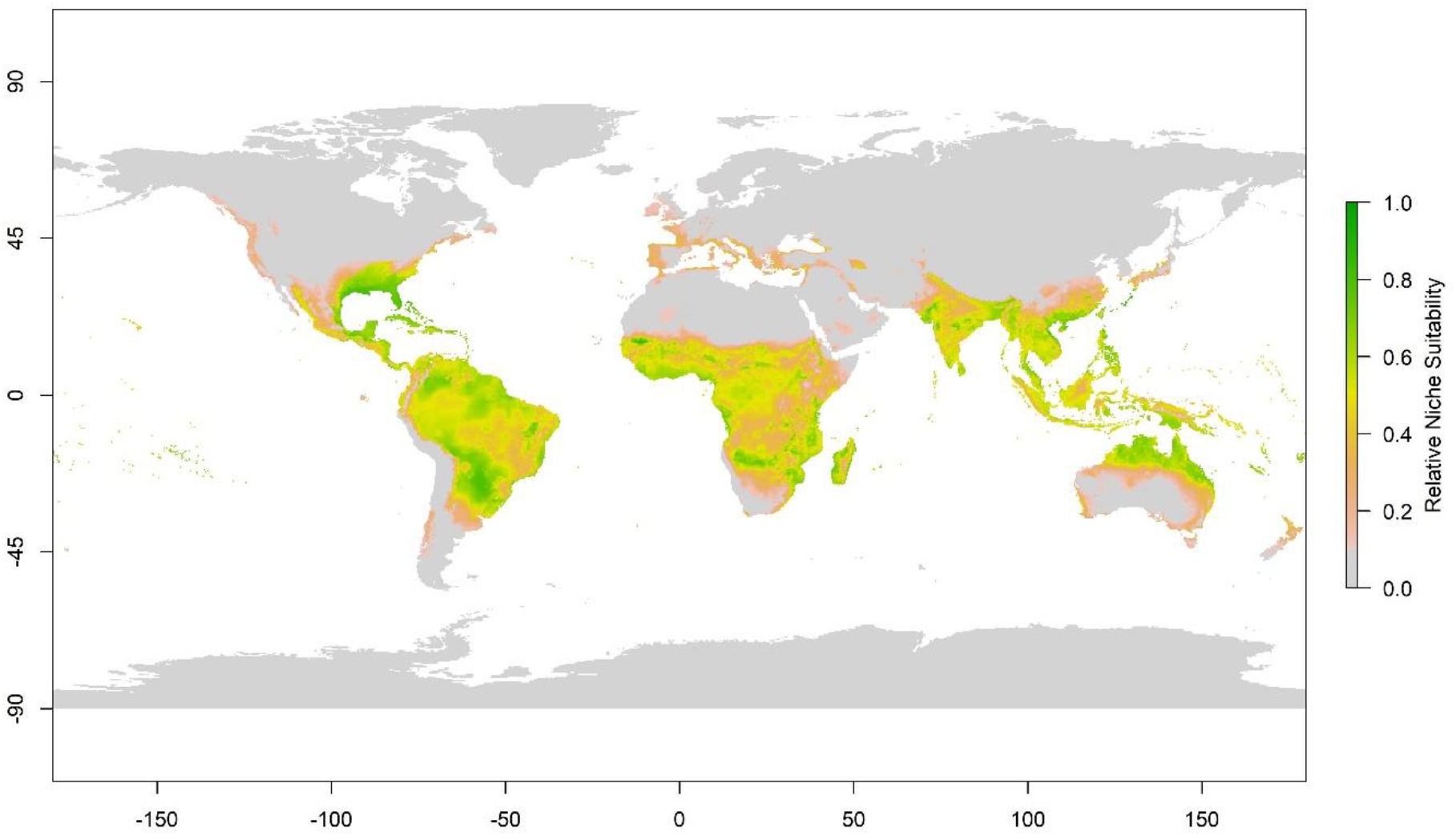
Map of relative mean niche suitability for modern crocodilians. Niche suitability was estimated using a MaxEnt semi-supervised classifier, trained using modern climate data and bootstrapped 50 times. Climate data was sampled using the geographic coordinates of crocodilian occurrences, together with synthetically generated pseudo-absence data.

**Fig 3.**
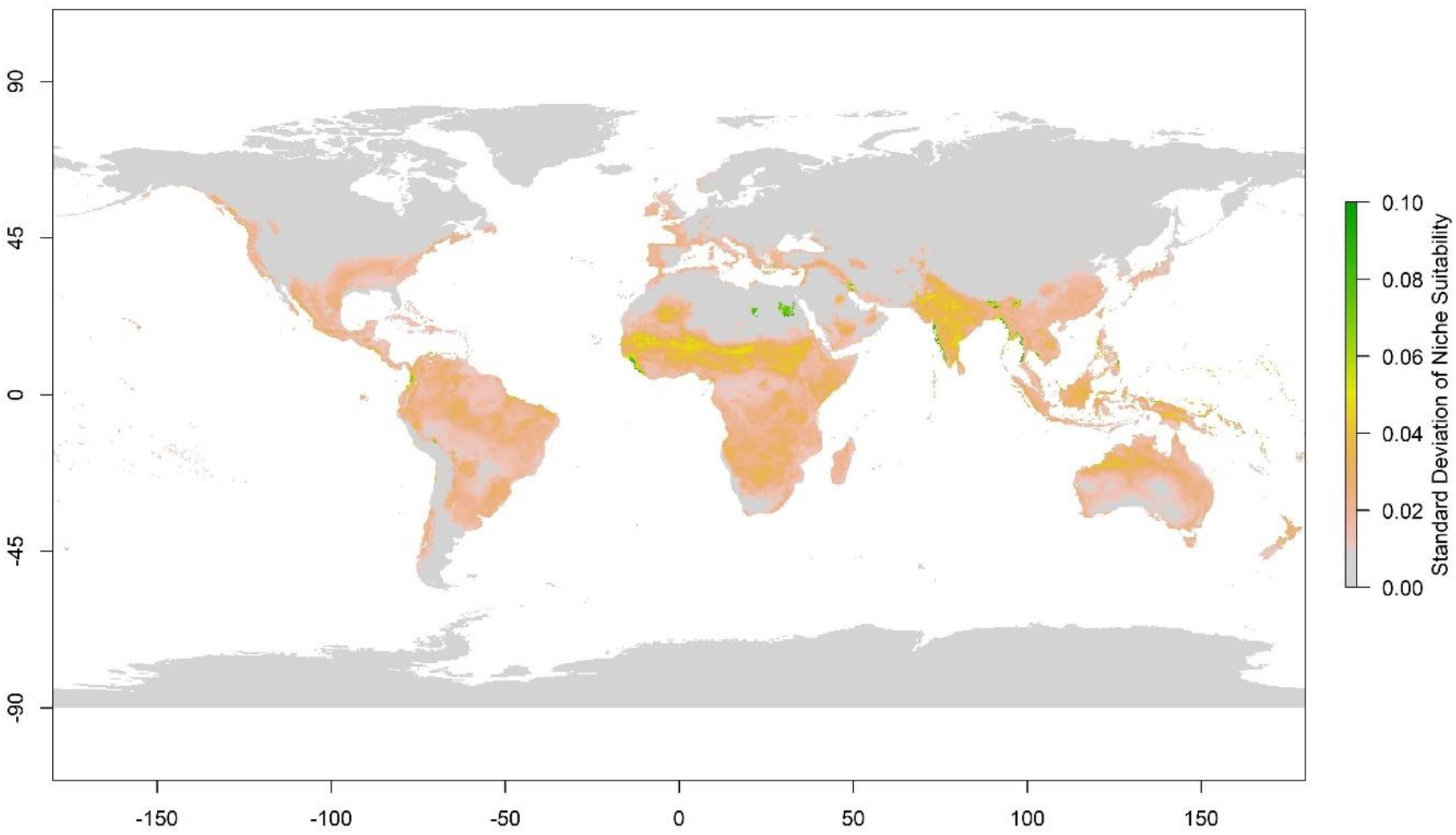
Map of standard deviation in estimated niche suitability across 50 bootstrap replicates. MaxEnt Models were each trained using a random 50% subsample of the complete dataset. Areas of highest model uncertainty are shown in areas where the average estimated niche suitability is low (Fig 2.).

There are a range of performance metrics that are applicable to maximum entropy classifiers (Elith et al., 2010[30]). This analysis uses the Area Under receiver Curve (AUC) metric, which expresses model performance as the plot area underneath a curve of true positives against false positives. This highest AUC possible is 1, a perfect classifier where 100% of the plot area is below the line and 100% of testing data is classified accurately. A classifier with an AUC of 0.5 would be classifying test data at random. A classifier with an AUC below 0.5 would be more likely to classify a test data point incorrectly than correctly [31].

The ENM model was used to predict the distribution of eusuchians in the Maastrichtian and Danian stages. These predictions were made by fitting the ENM to palaeoclimate variables derived from general circulation models simulating the Maastrichtian and Danian climate. The analysis used the same general circulation models used by Waterson et al [48] and Chiarenza et al [25]. These are derived from a lower-resolution variant of UK MetOffice model HadCM3, which is a fully-coupled atmosphere-ocean model. Climate variables were selected to match the bioclimatic variables used to train the ENM. These models have performed well in previous examples of deep-time ENM models [25, 48]. Validation efforts have raised questions about the efficacy of these models in continental interiors [61], so their inferences must but interpreted with caution. However, living crocodilians are associated with standing bodies of water [9] more likely to be associated with coastal and lowland areas than continental interiors. Similarly, fossil occurrences are expected to be associated with high levels of sedimentation, which may be more closely associated with continental margins.

The ENM was fitted the variables corresponding to those used to estimate the initial model. This yielded a map of niche suitability in the Maastrichitian and Danian stages. These maps of niche suitability were then thresholded using the same niche probability as the ENM of the present day.

Both maps of niche suitability and estimated geographic range were clipped fit a palaeogeographic map from Markwick [32]. Although there is debate about the accuracy of palaeogeographic reconstructions [33], both reconstructions have been used [32,33] and so their use here represents consistency across Eusuchia research.

The performance of the ENM when fitted to palaeoclimate data was then evaluated using fossil occurrences. Fossil occurrence data was downloaded from the Paleobiology Database (paleobiodb.org) on the 24^th^ of January 2021 using the search terms Eusuchia, Maastrichtian, and Danian (supplementary tables 3 & 4). Occurrence records were filtered to include only those between 72.1 and 66 Ma (Maastrichtian) and 66 – 61.6 Ma (Danian). Fossil entries associated with ootaxa, ichnotaxa, debris, and eggshell were removed to ensure only body fossils were included. Records with blank palaeo-coordinates and duplicate occurrences, identified by their occurrence number, were omitted. Palaeo-locations were plotted and visually checked for obvious anomalies. 12 Maastrichtian occurrences had no ID numbers and no further information and so were removed. Owing to the taxonomic uncertainty of the Eusuchia clade, a further search using the search terms Crocodyliforms and Neosuchia was undertaken to identify Eusuchia species that had been incorrectly assigned or misidentified. A species-level identification was used to verify their relationship to the Eusuchia clade. This yielded two additional Maastrichtian occurrences and no additional Danian occurrences. In total, 456 unique Maastrichtian and 126 Danian occurrences were used.

The ENM was subjected to further validation using the AUC performance metric. This validation was implemented using a combination of palaeoclimate data and fossil occurrences. The AUC scores of all three implementations are shown in table 1.

**Table 1:** AUC scores of MaxEnt model when validated using occurrence and climate data from 3 time intervals.

| **Model** | **AUC score** |
| --- | --- |
| Modern | 0.97 |
| Maastrichtian | 0.85 |
| Danian | 0.90 |

## Results

The Ecological Niche Model (Fig 2) of living eusuchians using contemporary climate data achieved an AUC score of 0.965 (Table 1). This indicates that the maximum entropy classifier correctly identifies suitable climate niches in 96% of the test data. This suggests that this species distribution model has a very high fidelity, and can be used to make accurate predictions when fitted to alternative climate scenarios.

Partitioning niche probabilities using a threshold value nominally divides a map into regions within a geographic range and regions outside it. The specificity and sensitivity of Ecological Niche Models is a function of this threshold, which is on the scale of the mapped niche probabilities. The specificity, or false positive rate, increases with the threshold value. The sensitivity, or true positive rate, is decreases with the threshold value. The threshold value used in this analysis was 0.565. This is the niche probability where the true positive and false positive rates are equal. Since niche probabilities lie on a scale from 0 to 1, this threshold value is close to the centre of the distribution. This suggests that any model errors are evenly distributed across both false positives and false negatives. When applied to the map of niche probabilities, this threshold predicts the geographic range of extant eusuchians is generally confined to regions within 35 degrees of the equator (Fig 4).

**Fig 4.**
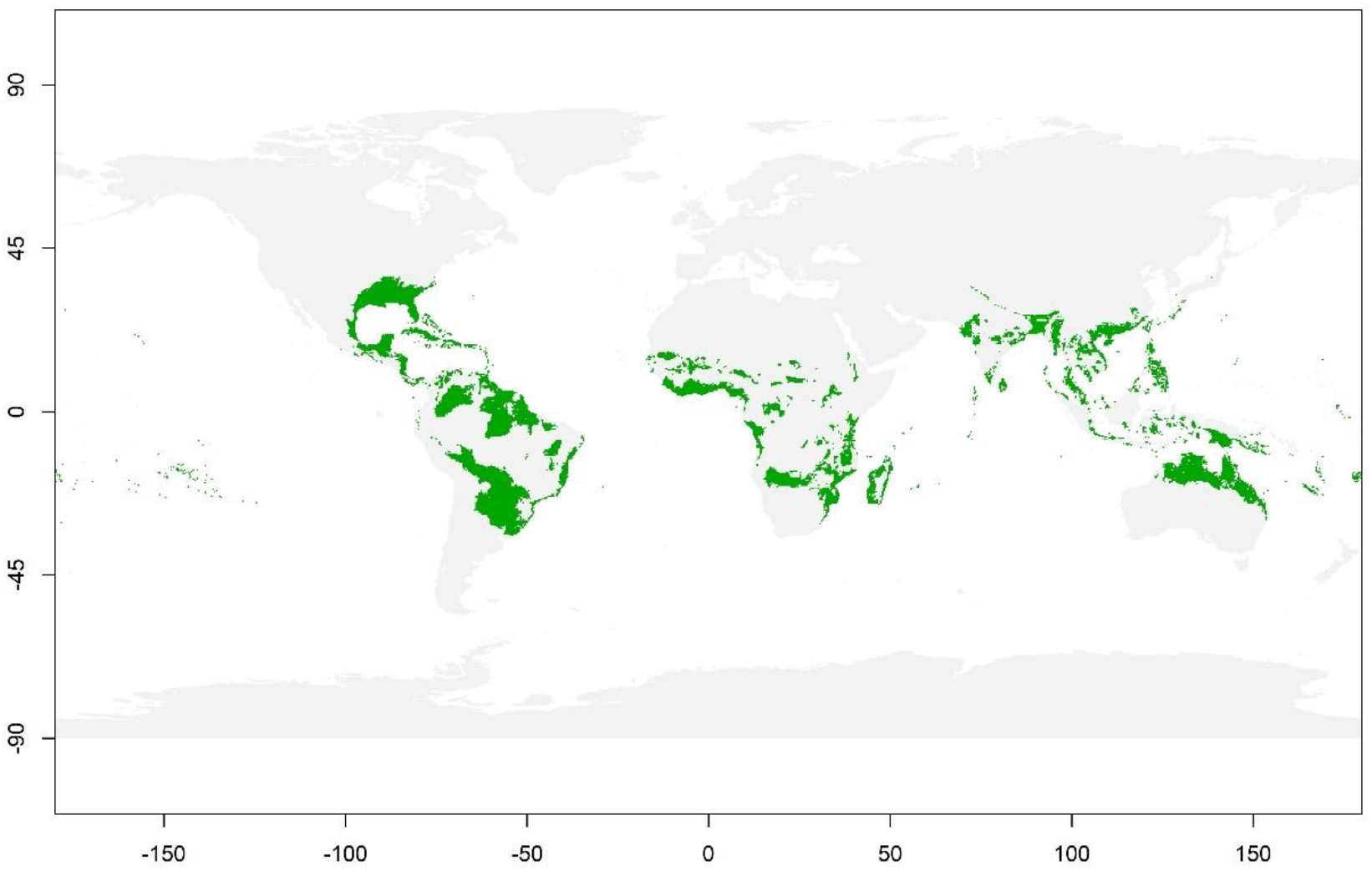
Map of predicted biogeography of living crocodilians. The cutoff threshold for niche suitability assumes an equal false-positive and false-negative rate of prediction error.

The standard deviation of the bootstrapped ENMs returned a map showing a generally even classifier certainty throughout the eusuchian’s predicted range. Climate niche uncertainty rarely exceeds 5% in any geographic region. Some isolated regions reach uncertainties of as high as 10%, however these regions all lie below the threshold value across all iterations and are not populated with observations.

The ENM returned relative contributions of each climate variable to the classifier. Precipitation in the wettest quarter contributed for the greatest variation, accounting for 50% of overall niche probability (Table 2). The second highest contributor was the mean temperature of the driest quarter, which accounted for 34% of niche probability. The temperature of the warmest quarter accounted for 7% of variation in niche probability. The remaining variables all accounted for less that 5% of variation (Table 2).

**Table 2:**
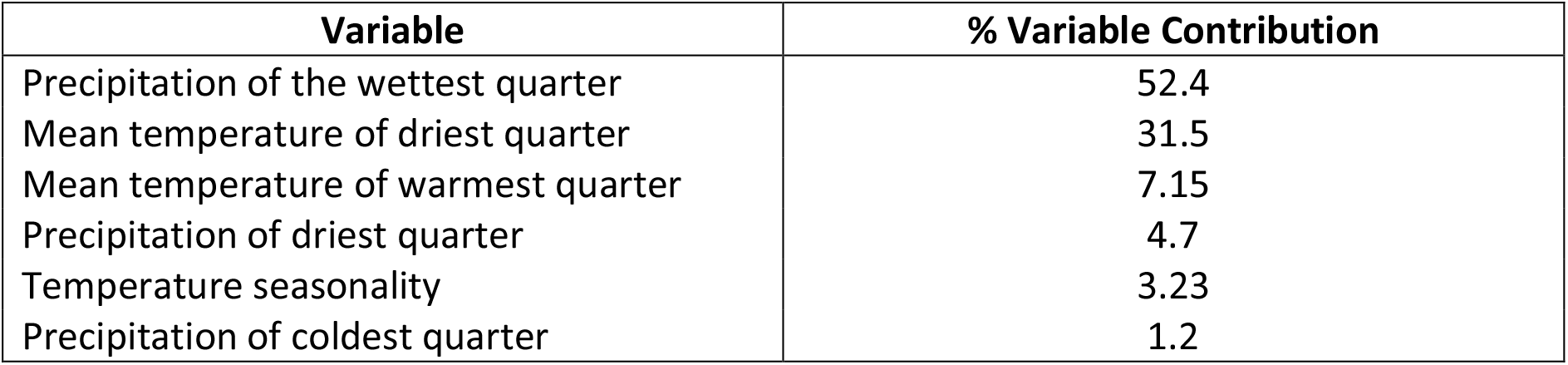
Percentage contributions of each climate variable in estimating niche suitability for crocodilians in a modern climate regime.

Response curves for each variable indicate that niche probability is highest in areas with approximately 1.5 meters of rainfall during the wettest quarter. Niche probability was highest in areas with a temperature of between 15°C and 20°C during the driest quarter, and areas above 30°C during the warmest quarter (Fig 5).

**Fig 5.**
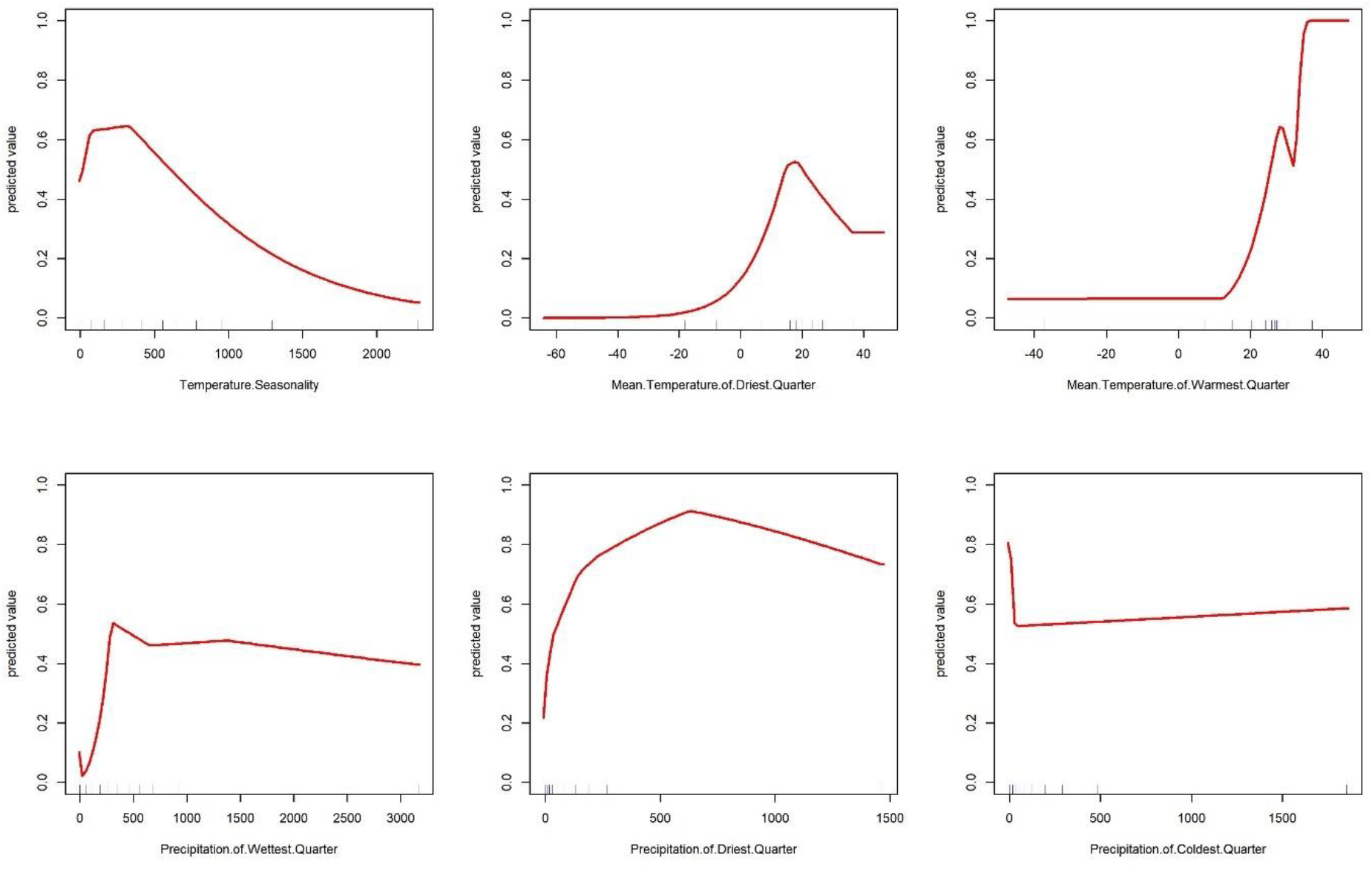
Response curves of each climate variable. Niche suitability is given on the y-axis for a given value for each variable.

When fitted to maps of palaeoclimate data, the ENM shows eusuchian niche probability to be associated negatively with distance from the equator (Fig 6 & 7). However, the latitudinal range of eusuchians is predicted to be very much wider than is predicted for the current climate. In the Maastrichtian, the latitudinal range of the eusuchians is predicted to be within 45 degrees of the equator (Fig 8). In the Danian, the latitudinal range of eusuchians increases to within 55 degrees of the equator (Fig 9).

**Fig 6.**
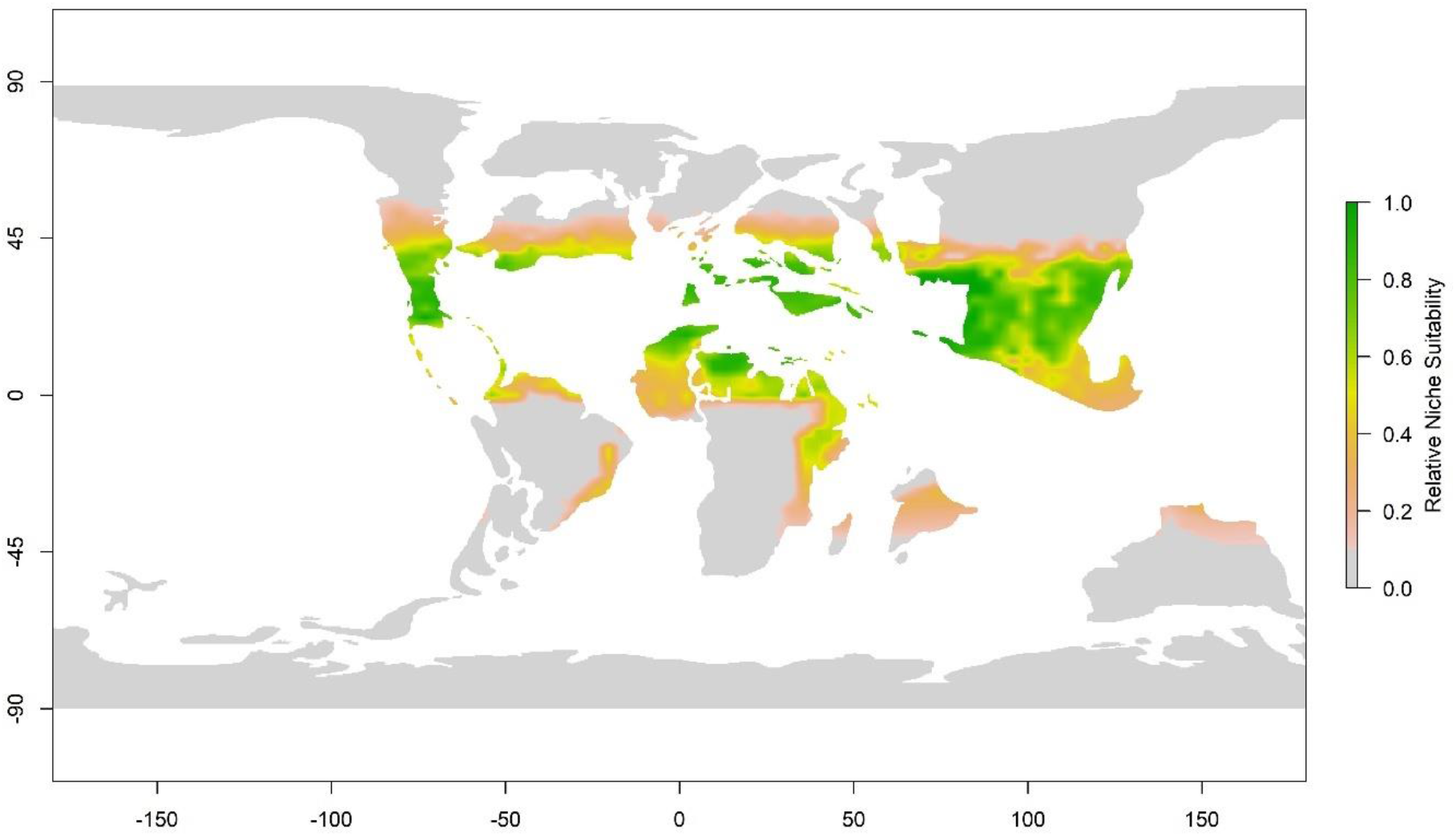
Map of predicted eusuchian niche suitability in the Maastrichtian stage of the Cretaceous period. A MaxEnt model trained using extant crocodilians and contemporary climate data was fitted to paleoclimate variables estimated using a general circulation model.

**Fig 7.**
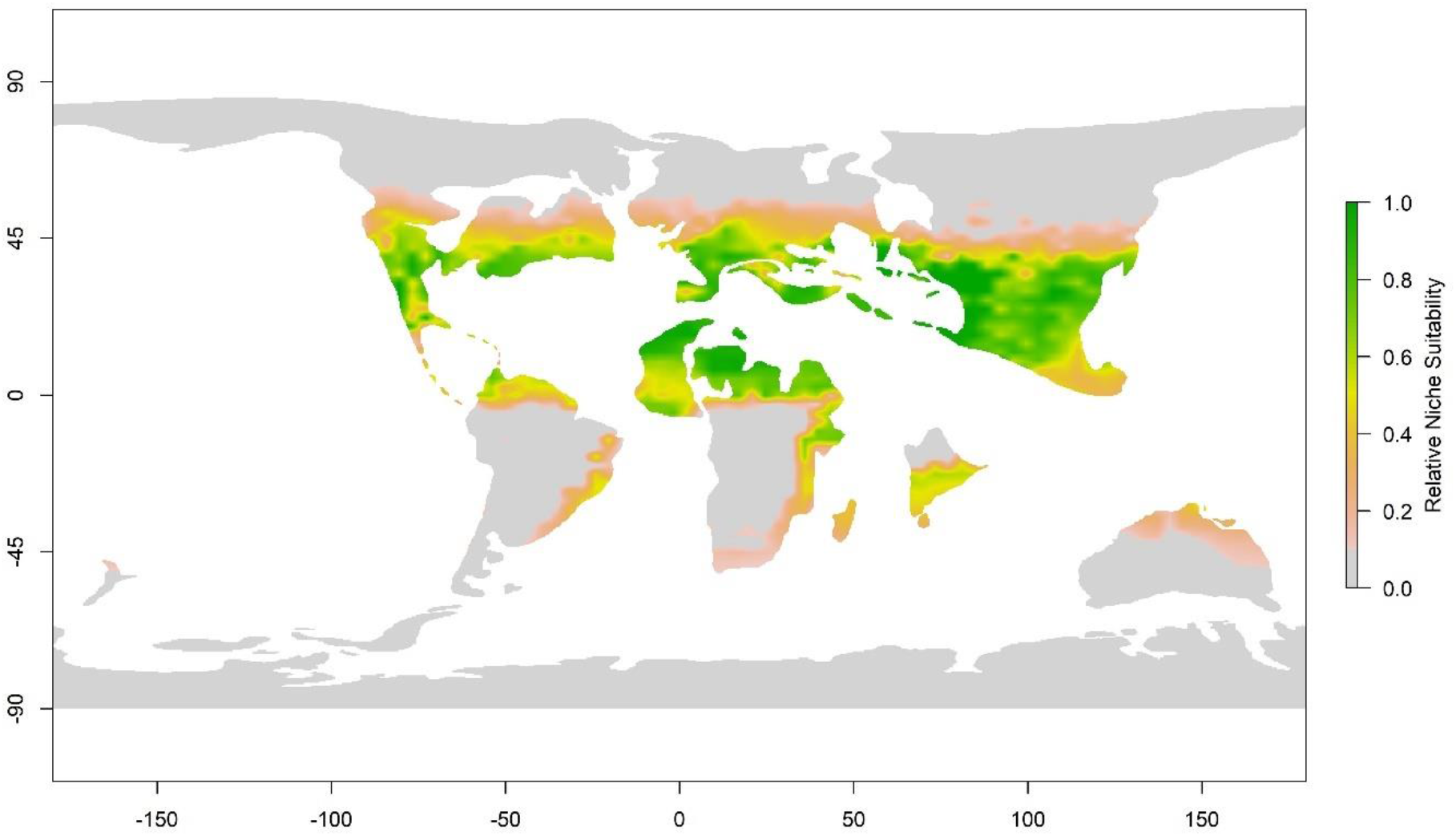
Map of predicted eusuchian niche suitability in the Danian stage of the Paleogene period. A MaxEnt model trained using extant crocodilians and contemporary climate data was fitted to paleoclimate variables estimated using a general circulation model.

**Fig 8.**
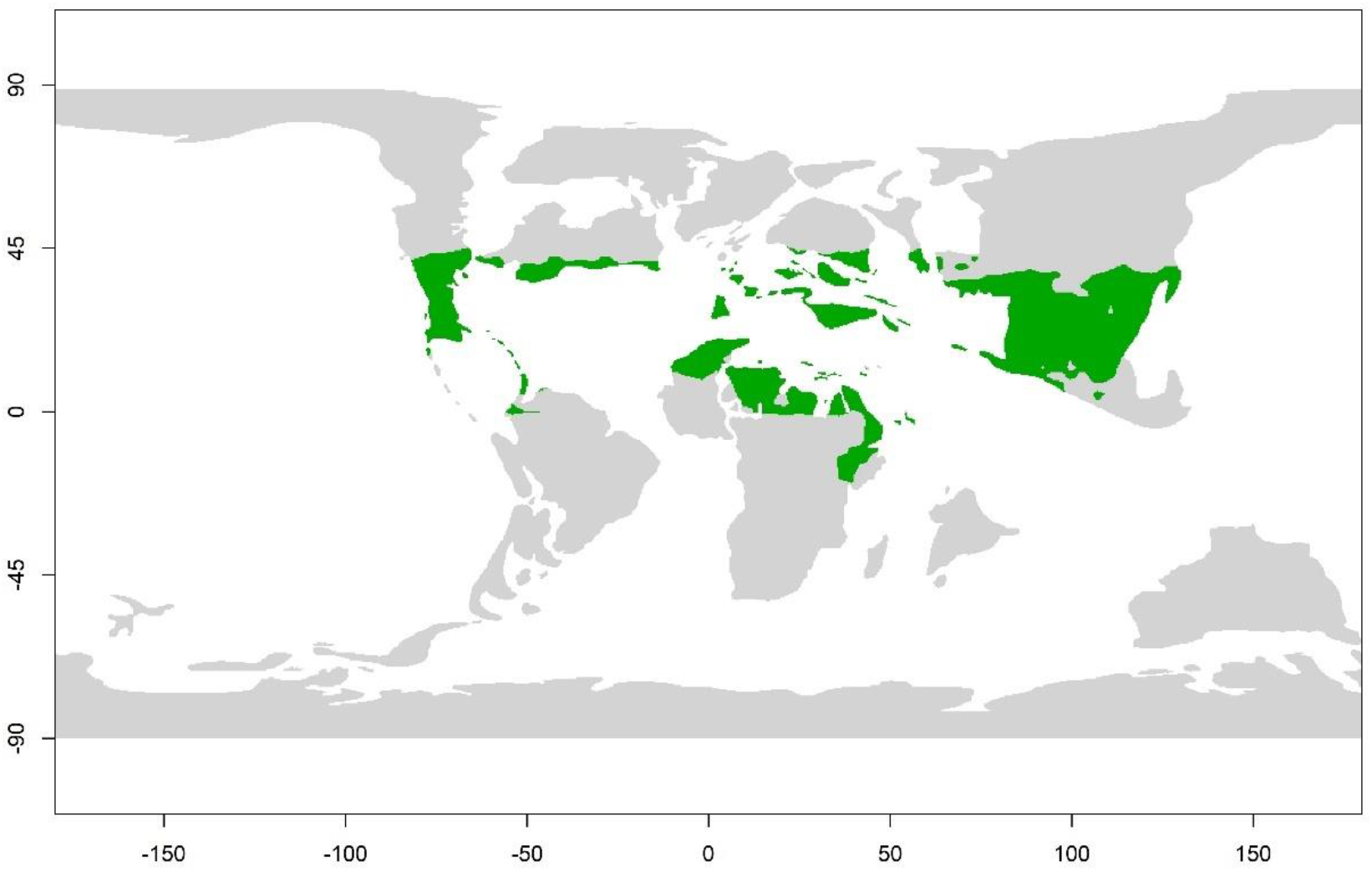
Map of predicted realised niche for eusuchians in the Maastrichtian stage of the Late Cretaceous period. Maastrichtian niche probability (Fig 6.) is thresholded using the same value as the contemporary model (Fig 4.), which assumes an equal false-positive and falsenegative model error rate.

**Fig 9.**
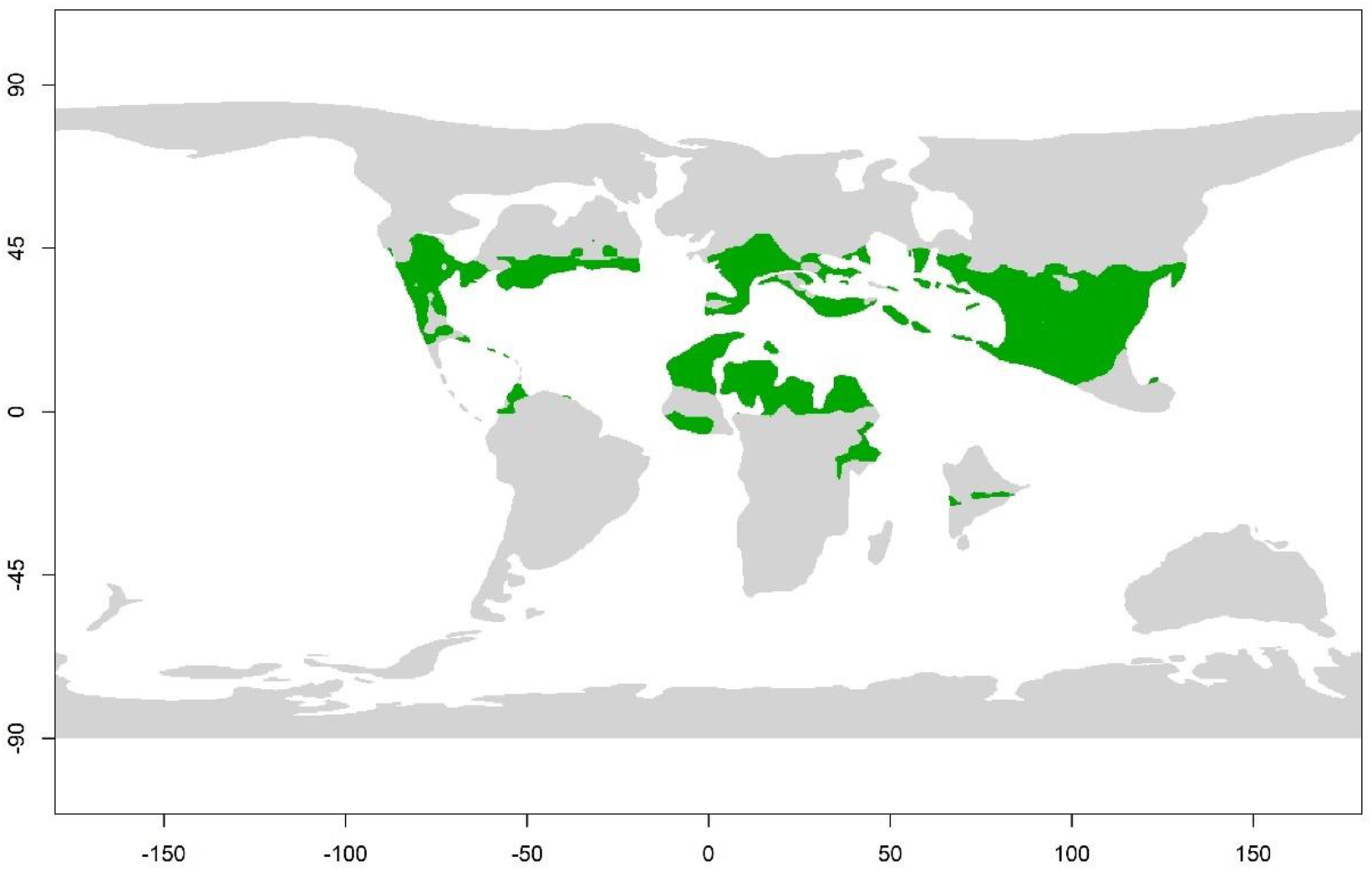
Map of predicted realised niche for eusuchians in the Danian stage of the Paleogene period. Danian niche probability (Fig 6.) is thresholded using the same value as the contemporary model (Fig 4.), which assumes an equal false-positive and false-negative model error rate.

The ENM classifier performs well in predicting fossil palaeogeography. Fossil occurrence data and maps of palaeoclimate variables return favourable AUC scores of 85% accuracy for the Maastrichtian stage and 90% for the Danian stage (Table 2). The decrease in classifier accuracy may be attributable to the smaller sample size of fossil occurrences, or the lower spatial resolution of the palaeoclimate data.

## Discussion

The results of the analysis confirm a close relationship between the geography of extant eusuchians and the climate, as has been established in previous studies[9]. The role of temperature in the geographic distribution of modern crocodilians seems likely to be due to their ectothermic physiology. The inability of crocodilians to generate their own body heat makes them dependent on environmental conditions to sustain their metabolic processes, and therefore confining them warmer regions. Markwick [9] postulated that a coldest month minimum temperature of 5.5 °C represents a climatic limit on Eusuchia niche suitability, and the results of the ENM analysis do not contradict this. Eusuchia niche suitability does not decrease at temperatures above 38 °C, suggesting that modern environmental temperatures do not represent the upper thermal limit of the Eusuchia clade. This mitigates the concerns raised by Rogers et al [34] and Sunday et al [35] who suggest that extant Eusuchia are approaching their upper thermal limits.

Perhaps surprisingly, precipitation ostensibly has greater significance in constraining the biogeography of eusuchians than temperature. The exact mechanism for this is deserving of further study. It is intuitive that some regions are too dry for eusuchians to survive. Their amphibious ecomorphology would seem to limit them to areas with sufficient rainfall to sustain fluvial, lacustrine or coastal environments. Another possible factor may be the role of humidity in the successful incubation of eusuchian eggs [36]. Precipitation may also interact with eusuchian physiology. Water changes temperature more slowly than air, and so can cushion aquatic organisms from rapid temperature changes. Therefore, an aquatic ectothermic organism may remain more active active during periods of decreased temperature. This is reflected in the behaviour of living crocodilians, which are often more active at night [1]. Such possible interaction between the biogeography of eusuchians and precipitation corroborate previous findings linking crocodilian distribution to bodies of standing water[9].

The close relationship between known occurrences and predicted niche suitability suggests that the fundamental and realised niche of extant Eusuchia is broadly similar. This supports the findings of Morales-Betancourt et al [37] and Balaguera-Reina et al [38], who constrained the niche suitability of *Crocodylus intermedius* and showed a similar intimacy between the predicted and fundamental niche, albeit at a regional scale.

There are some areas where niche suitability is ostensibly high, however there are no occurrences from these regions included in the training data. This may be due to a lack of sampling in these regions, or a genuine absence of crocodilians from these regions for reasons other than climate. This may include persecution by humans, development of coastal habitats by humans, and unsuitable topography. It is possible a co-dependent organism, such as a prey item, may be dictated to a greater extent by climate than are eusuchians. In this scenario, the constraints upon eusuchian biogeography by climate would be only secondary. However, modern crocodilians have an extremely broad diet including hypercarnivory, piscivory, insectivory, durophagy and scavenging [1]; it seems unlikely that their geographic range would be limited indirectly by other organisms.

The ENM estimated in this analysis demonstrates that climate variables are highly effective in predicting the biogeography of fossil eusuchians. This has implications for the physiology of extinct eusuchian species, due to the apparent relationship between environmental conditions than the metabolic rates of modern crocodiles. It seems clear that extinct members of the crocodile crown-group had a similar physiology to their modern-day counterparts. This further implies that ectothermic physiology is a synapomorphy of the crown-group crocodilians, and that ectothermy has not arisen convergently in modern species. Members of the Eusuchia do occur prior to the common ancestor of modern crocodiles [39], and the results of this ENM analysis suggest that ectothermic physiology predates the crocodile crown-group. Recent speculation has suggested the crocodile lineage, the Pseudosuchia, are descended from an endothermic ancestor, and this condition was lost in more modern crocodylomorphs [40]. The results from this ENM analysis are consistent with previous work which concluded that endothermy had been lost prior to the evolution of the Mesoeucrocodylia, a higher-order clade which includes the Eusuchia [41].

The wider predicted latitudinal range of eusuchians during the Maastrichtian and Danian stages concurs with previous analysis by Markwick [2], who proposed a geographic distribution of between ~60°N and ~60°S. This increased range can be attributed to higher temperatures and higher rates of precipitation during. Such favourable climate conditions would have enabled eusuchians to colonise regions as far north as Canada and Europe. The ecospace available to eusuchians was greater in absolute terms. However, in practice this area of available ecospace would have been interrupted by geographic obstacles such as oceans. Such geographic obstacles would create conditions for genetic isolation, and therefore allopatric speciation. Therefore, the favourable climate conditions during the Maastrichtian and Danian stages may contribute to the high species richness of fossil eusuchians.

The predicted geographic distribution of Eusuchians during the Danian extends to a greater latitudinal range than in the Maastrichtian. This tracks the increase in global temperatures at the start of the Cenozoic [42]. This suggests that climate conditions overall were becoming more favourable for eusuchians, and so any loss in diversity during the Cretaceous-Palaeogene boundary cannot be attributed to climate change. Eusuchian extinctions during the Cretaceous-Palaeogene boundary are therefore more attributable to the end-Cretaceous bolide strike and its associated impacts [43]. The decline in eusuchian diversity during the Cenozoic [8] was either not initially attributable to climate change, or that decline in diversity began some time after the Danian stage. Further, climate seems not to be attributable to the loss of ecomorphological diversity across the Cretaceous-Palaeogene boundary [12].

All living crocodilians exhibit amphibious behaviour. As noted previously, living in water may serve to cushion them from temperature changes. Therefore, such amphibious behaviour may interact with physiology in constraining suitable conditions for eusuchians. This may limit the utility of the ENM in this analysis to eusuchians which exhibit amphibious behaviour. None of the eusuchian taxa included in this study have ostensibly terrestrial features, which may contribute to the relatively high AUC scores reported here. However, some fossil eusuchians exhibit a secondarily terrestrial morphology, for example the Planocraniidae [44] and the Mekosuchinae [45]. Members from these families are not known from the Danian, but it is plausible that a secondarily terrestrial ancestor may have existed. It is deserving of note that the ENM synthesized in this analysis may be limited only to amphibious forms, and that secondarily terrestrial Eusuchia may have occurred outside of the predicted geographic range. Similarly, secondarily terrestrial Eusuchia may also have had a more limited geographic range within the area predicted for the group as a whole.

The distribution of eusuchians may not have been homogeneous within the geographic range predicted by the ENM. Some areas suitable for eusuchians may have been empty of examples due to factors other than climate, for example geographic isolation or the presence of competitors. Eusuchian subtaxa may also have been adapted to more limited climate niches within the predicted range of the group as a whole. Ecological niche modelling of eusuchian subclades, such as the Alligatoridae or Crocodylidae, would be an interesting avenue of further research. However, such analyses would be constrained by much smaller sample sizes and would be difficult to validate.

The analysis presented here confirms the fundamental role of climate in determining the biogeography of both living and fossil crocodiles. Temperature is of great importance, as suggested by previous work, however the role of precipitation is even greater. During the Maastrichtian and Danian stages, eusuchians occupied a much greater latitudinal range than their extant counterparts. The fundamental niche available to eusuchians increases across the Cretaceous-Palaeogene boundary, suggesting that climate was not a limiting factor for eusuchian diversity at this time. The performance of climate variables in predicting the distribution of crocodile fossils suggests that fossil eusuchians were physiologically similar to modern forms, and that ectothermy is a synapomorphy of the crocodile crown-group.

